# Comparison of Principal Component Analysis and t-Stochastic Neighbor Embedding with Distance Metric Modifications for Single-cell RNA-sequencing Data Analysis

**DOI:** 10.1101/102780

**Authors:** Haejoon (Ellen) Kwon, Jean Fan, Peter Kharchenko

## Abstract

Recent developments in technological tools such as next generation sequencing along with peaking interest in the study of single cells has enabled single-cell RNA-sequencing, in which whole transcriptomes are analyzed on a single-cell level. Studies, however, have been hindered by the ability to effectively analyze these single cell RNA-seq datasets, due to the high-dimensional nature and intrinsic noise in the data. While many techniques have been introduced to reduce dimensionality of such data for visualization and subpopulation identification, the utility to identify new cellular subtypes in a reliable and robust manner remains unclear. Here, we compare dimensionality reduction visualization methods including principle component analysis and t-stochastic neighbor embedding along with various distance metric modifications to visualize single-cell RNA-seq datasets, and assess their performance in identifying known cellular subtypes. Our results suggest that selecting variable genes prior to analysis on single-cell RNA-seq data is vital to yield reliable classification, and that when variable genes are used, the choice of distance metric modification does not particularly influence the quality of classification. Still, in order to take advantage of all the gene expression information, alternative methods must be used for a reliable classification.

## Introduction

Improving sequencing technologies in the past several decades has enabled single-cell sequencing to shed light on the understanding of cellular relationships within higher organisms^1^. In the past several years, studies focusing on single cell sequencing of whole transcriptomes have already revolutionized our understanding of the complexity of eukaryotic transcriptomes^1–3^. Single cell RNA sequencing allows us to explain the phenotypic heterogeneity^4^ observed in certain groups of cells and tissues by profiling cell-to-cell transcriptional heterogeneity. A major application of single-cell RNA-seq results in its ability to identify rare (novel) subtypes^3,5–9^ that are not easily identifiable using pre-known factors^1^. For example, single cell RNA sequencing has been exploited to identify novel subtypes in the placenta^6^ and intestine^5^. Single cell RNA sequencing therefore has great implications in studies regarding embryonic developments^6^, adult stem cells^1^ and cancer^10^ as it allow us to investigate and understand the heterogeneity in cell populations, as well as find novel subtypes.

Although single-cell RNA-seq has already enabled recapitulation of known cellular subtypes^3,5^, additional computational methods are needed to identify and characterize new subtypes^1^. This task is often complicated by multiple factors such as the high dimensionality of the data, accompanying intrinsic noise, drop-outs during library preparation, and the data’s stochastic nature^11^. In visualizing the data, however, a plethora of methods have already been developed in attempt to visualize single-cell RNA-seq data in order to visually identify subtypes^12^ with dimension reduction. An often extensively used tool is Principal Component Analysis (PCA) (11), as well as the default Euclidean distance metric for t-Stochastic Neighbor Embedding (tSNE). PCA, however is a linear transformation of data, and the default Euclidean distance has been previously known to work poorly on high dimensional data^13^. Therefore, we hypothesized that alternative distance metrics aside from the commonly used PCA and default Euclidean for tSNE will be better in classifying subtypes.

Here, we apply and assess the performance of multiple alternative distance metrics to a publicly available single-cell RNA-seq dataset^9^, whose subtypes were annotated^9^. The metric modifications for tSNE and PCA were assessed in terms of their ability to classify subtypes reliably, and robustness.

## Methods

### Processing of previously published single-cell RNA-seq data

SRA files for each study were downloaded from the Sequence Read Archive (http://www.ncbi.nlm.nih.gov/sra) and converted to FASTQ format using the SRA toolkit (v2.3.5). FASTQ files were aligned to the human reference genome (hg19) using Tophat (v2.0.10) with Bowtie2 (v2.1.0) and Samtools (v0.1.19). Gene expression counts were quantified using HTSeq (v0.5.4).

Linnarson data, a single-cell RNA-seq data for cell types in the mouse cortex and hippocampus^9^ (Zeisel, 2015 #2) was used in R, and matched with corresponding tags for previously known subtypes. Those with unknown or tag less subtypes were omitted.

The data was log transformed and scaled before having performed any kind of distance metric quantification.

## Principal component analysis and tSNE

For all pairs of the previously known subtypes specified^9^, principal component analysis, and t-Distributed Stochastic Neighbor Embedding (t-SNE) with various distance metrics embedded, were performed. Linear Discriminant Analysis (LDA) from MASS (v7.3 – 45) was used as a classifier for each pair of subtypes. For principal component analysis, prcomp from R stats package v(3.2.2) was used then visualized into a scatterplot including variance, with the first two principal components. tSNE was performed using Rtsne (v0.11) with a specific distance metrics assigned (Euclidean, Manhattan, Minkowski).

For both PCA and tSNE, the classification was then evaluated and visualized using ROCR 1.0-5.gz with an AUC value and ROC curve plot.

### Performance Benchmarking

To assess the robustness of PCA and tSNE with different distance metrics, we sought to benchmark performance as a function of the number of genes used in the dimensionality reduction. Varying number of genes was randomly sampled on a logarithmic scale. Performance AUC for PCA and tSNE with distance metrics embedded were calculated as noted, then replicated ten times with different seed values. The AUC values for all metrics were obtained from calculating the performance values of predictions for LDA. Resulting matrices of difference metrics with replications and gene sizes were melted with reshape2 (v1.4.1). Geom_smooth from ggplot2 (v1.0.1) was then used with loess to create the final benchmarking plot containing the mean lines of benchmarking performance AUC values for PCA and tSNE with distance metrics.

From visual inspection of scatterplots including ROC curve and AUC value of all possible pairs of subtypes from previous runs (PCA and tSNE), we observed that the pair Pyramidal SS and Pyramidal CA1 was the most similar and hence most challenging to differentiate, so further runs were focused on this pair. In addition, we benchmarked performance using the 200 genes with the most variance in gene expression across cells. Within these variable genes, gene samples were randomly selected again on a logarithmic scale to create final benchmarking plots.

## Result and Discussion

When trying to identify the subtypes of a data whose subtypes are not annotated, many researchers use PCA or tSNE with default Euclidean metric for single cell RNA sequencing data (Figure 1), and visually identify the different subtypes. The visualized plot contains clusters of points, which are often distinguished into groups or subtypes depending on the approximate distance between the clusters. Therefore, to quantify how well one subtype is classified from another when the choice of distance metrics is varied, all possible pairs of subtypes annotated were tested (Figure 2), for different metrics. The scatterplots of all subtype pairs for PCA and tSNE with Euclidean were mostly visually well classified with AUC values near 1 (perfect classification) except Pyramidal SS and Pyramidal CA1 (Figure 2). Further tests with various distance metrics were therefore run on the Pyramidal CA1 and Pyramidal SS pair as our aim was to observe which metric performed better when the classification became more difficult (when subtypes are more similar).

**Figure 1:**
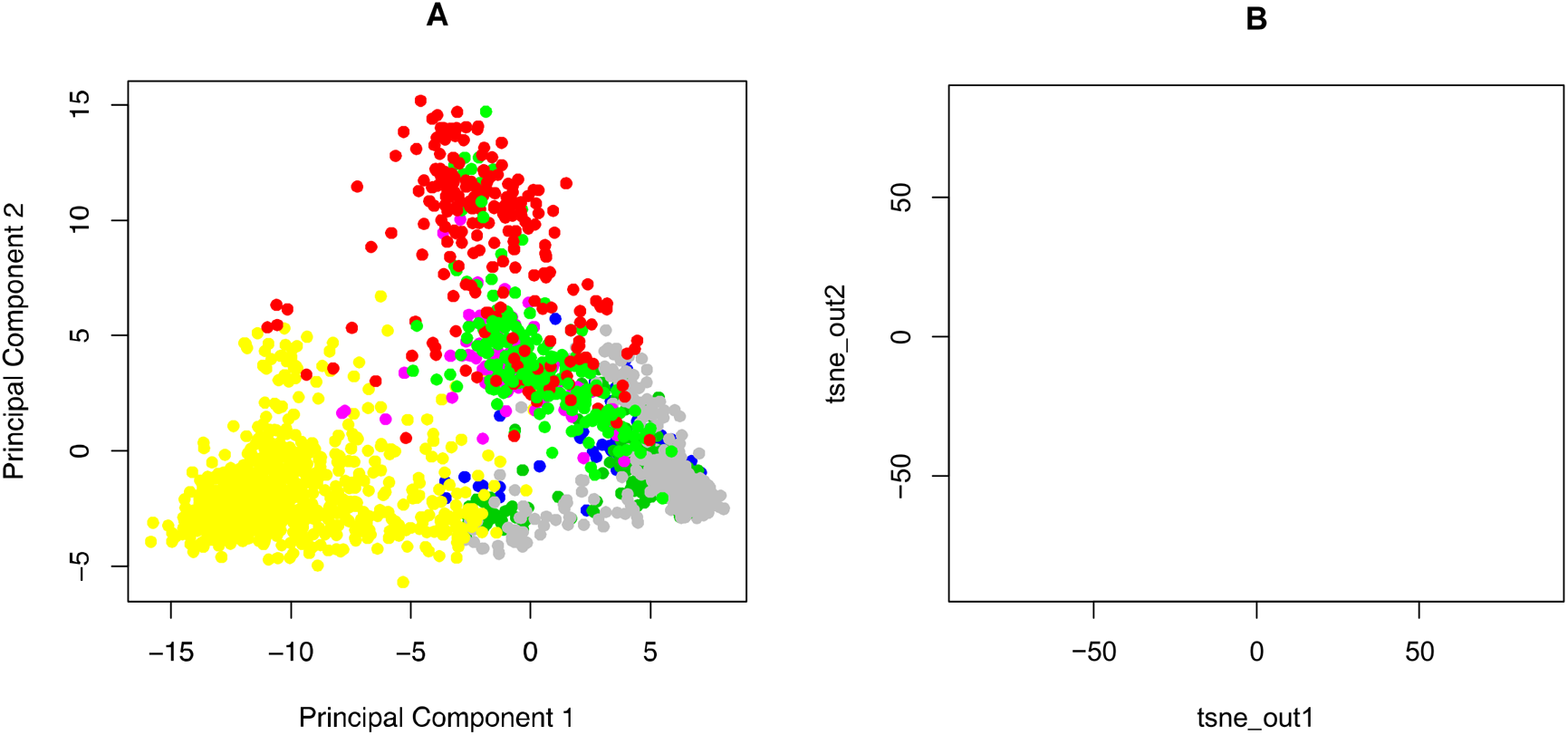
Dimensionality reduction and visualization of single cell RNA-seq data from Linnarson et al.^9^ highlights limited ability to visually distinguish annotated subtypes. a. Dimensionality reduction and visualization using PCA. 100 genes with the greatest variance post log transformation (See Methods) were selected and analyzed using PCA, then visualized using the first two principal components. The seven major subtypes annotated in the original paper are colored. We can visually distinguish the points into groups or subtypes depending on the approximate distance between the clusters.
b. Dimensionality reduction and visualization using tSNE. Again, 100 genes with the greatest variance post log transformation (See Methods) were selected and analyzed with tSNE using the default Euclidean distance metric, and colored with the annotated subtypes. Similar to the PCA plot, we can approximately distinguish some clusters.

**Figure 2;.**
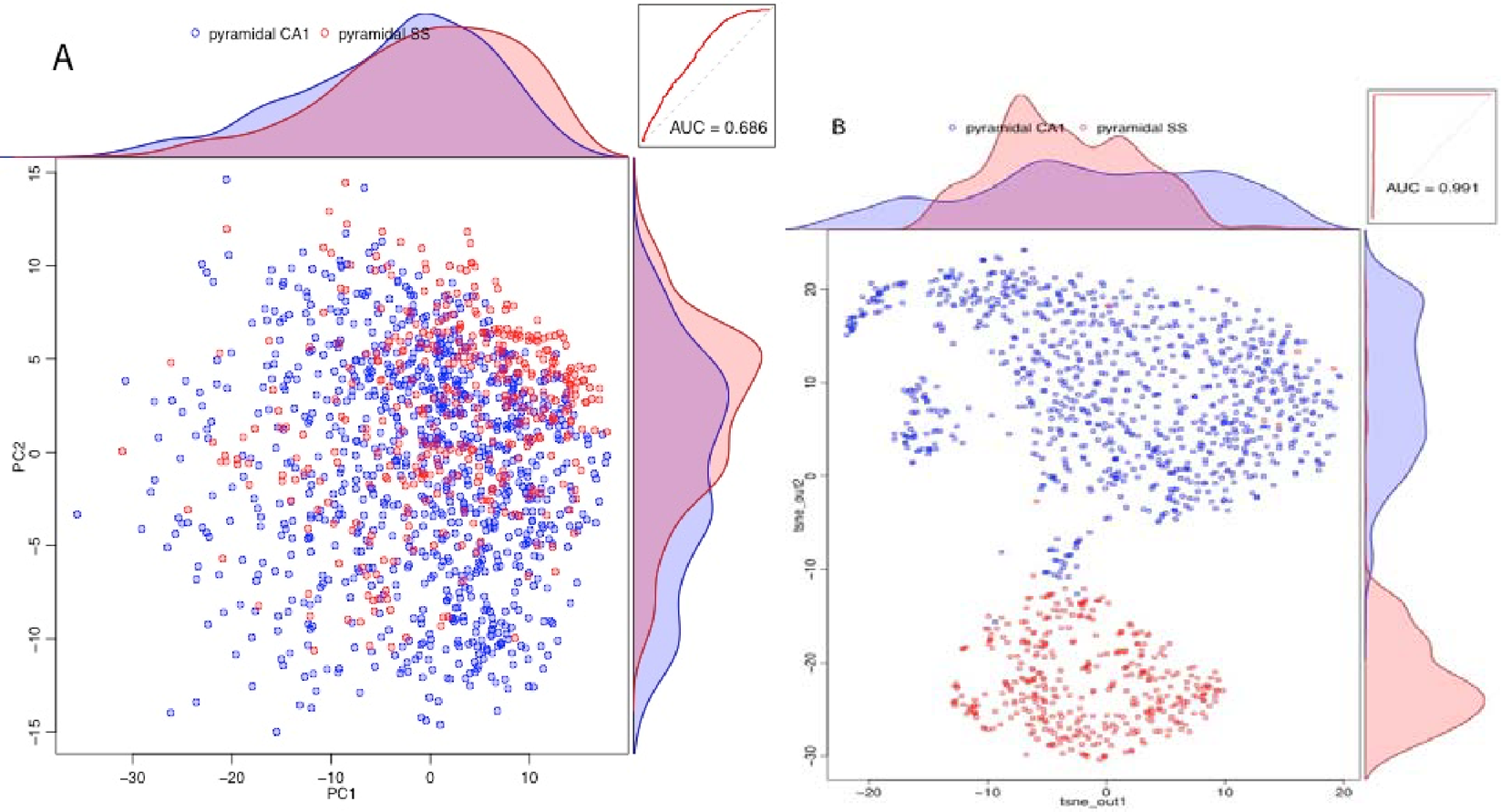
Quantification of subtype discrimination by LDA, ROC, and AUC. a. Distinguishing subtypes using PCA. All possible pairs of the major subtypes including astrocytes ependymal, endothelial-mural, interneurons, microglia, oligodendrocytes, pyramidal CA1, and pyramidal SS were run with PCA and visualized using a scatterplot. An LDA classifier (See Methods) was used to quantify the extent to which the subtypes can be distinguished and classified. ROC curves and AUC value were used for quantification of discrimination performance. All the genes were used. We observed that all the subtype pairs did nearly equally well with high AUC values, except for Pyramidal SS and Pyramidal CA1 as shown. Further tests with various distance metrics were therefore run on the Pyramidal CA1 and Pyramidal SS pair as our aim was to observe which metric performed better when the classification became more difficult (when subtypes are more similar).
b. Distinguishing subtypes using tSNE with Euclidean distance. 100 most variable genes were selected for the tSNE analysis. We observed that tSNE with Euclidean performed better than PCA as the AUC performance values were much higher compared to that of PCA for the same subtype pair.

In attempt to compare the performance of distance metrics, the trend in the performance of each metric over various sample sizes on a logarithmic scale, were observed. Lowering the size of the gene sample provides less information for the classification, so while the performance values may be lower with small sample sizes, a superior distance metric is expected to attain high and consistent performance values even with a slight increase in the sample size.

Initially, gene samples were selected randomly and benchmarked for multiple distance metrics (Figure 3, 1). There is a positive correlation between sample size and performance values across all metrics, but the performance values do not near 1, with 1 indicating complete separation of subtypes. This is important, as it is difficult compare to conclude which metric performs better when the performance values of all the metrics are similarly poor. Moreover, PCA and tSNE with Minkowski (min) and Manhattan (man) contain plateaus in their paths as sample size increases, suggesting that regardless of the number of genes sampled, the performance values is not likely to reach 1. We hypothesize this is due to the intrinsic noise in the single cell RNA sequencing data that complicates the classification as sample size increases. To resolve this issue, we took the 200 genes with the greatest variance in their gene expression and benchmarked within these variable genes (Fig3, 2). 200 genes were enough to allow the performance AUC values to reach 1 and to eliminate the plateaus. All distance metrics showed the characteristics of a superior distance metric, as performance values rapidly increased as sample gene size increased, and remain consistent.

**Figure 3:**
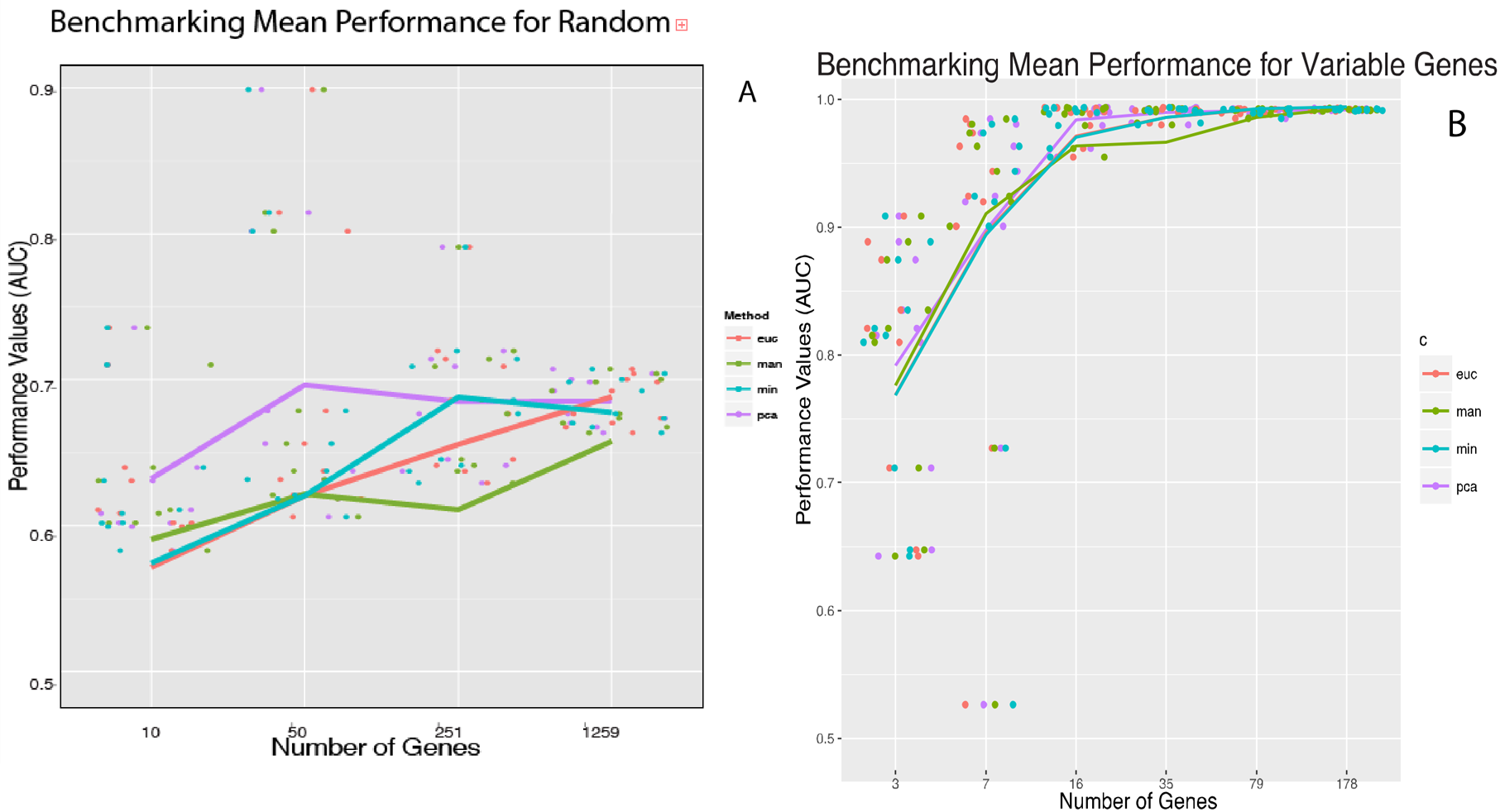
Benchmarking mean subtype discrimination performance for random and variable gene samples shows improved performance for variable gene samples. a. Benchmarking performance for random gene samples. In attempt to compare the performance of distance metrics, the trend in the performance of each metric over various gene sample sizes on a logarithmic scale, were observed. Lowering the size of the gene sample provides less information for the classification, so while the performance values may be lower with small sample sizes, a superior distance metric is expected to attain high and consistent performance values even with a slight increase in the sample size. Gene samples were selected randomly and benchmarked for multiple distance metrics. There is a positive correlation between sample size and performance values across all metrics, but the performance values do not near 1 for any metrics, suggesting imperfect classification even when high number of genes is used. Moreover, PCA and tSNE with Minkowski (min) and Manhattan (man) contain plateaus in their paths as sample size increases.
b. Benchmarking performance for variable gene samples. We hypothesize that the plateau and low AUC values were due to the intrinsic noise in the single cell RNA sequencing data that complicates the classification as sample size increases. To resolve this issue, we took the 200 genes with the greatest variance in their gene expression and benchmarked within these variable genes. 200 genes were enough to allow the performance AUC values to reach 1, as well as eliminate the plateaus. All distance metrics showed the characteristics of a superior distance metric, as performance values rapidly increased as sample gene size increased, and remain consistent, suggesting similar robustness as well.

We conclude that when using single cell RNA sequencing data, selecting the variable genes and omitting the noise in the data before analysis drastically improves the ability to classify subtypes. When the variable genes are selected, the subtype classification performance remains high regardless of method. If all genes are used, however, we find that an alternative, more sensitive, and robust method may be necessary to reliably identify subtypes. Therefore, while our findings may help those analyzing single-cell RNA-seq data for subtype classification and understanding heterogeneity, identification of rare, novel subtypes will require a more robust method.

